# Carbohydrate Metabolism Differs in Infants by Asthma-risk Status and is Associated with the Functional Potential of *Bacteroides cellulosilyticus*

**DOI:** 10.64898/2026.04.28.721144

**Authors:** Holly M. Steininger, Carlos E. Iglesias-Aguirre, Ariane R. Panzer, Juliana Durack, Michelle McKean, Michael D. Cabana, Spencer Diamond, Susan V. Lynch

## Abstract

2.

Childhood atopic disease is linked to delayed gut microbiome development and metabolic dysfunction, however microbial drivers remain unclear. To explore microbial correlates of asthma risk during a time of active gut microbiome development, we analyzed stool from 6-month-old infants at high asthma risk (HR) or healthy controls (HC), using Genome-resolved metagenomics (HR=7; HC=12) and untargeted metabolomics (HR=11; HC=15). We recovered 82 bacterial species-level metagenomic-assembled genomes (MAGs). Global Taxonomic composition did not differ by asthma risk. Anticipating that key differences might associate with specific genomes, a machine-learning approach pinpointed *Bacteroides cellulosilyticus, Hungatella effluvii*, and *Enterocloster aldenensis* as linked with asthma risk status. All three species were more abundant in HC infants and the *B. cellulosilyticus* genome was enriched for carbohydrate metabolism genes relative to other MAGs. Metabolomic profiling revealed variance associated with asthma risk (PERMANOVA, R^2^ =0.069, p=0.016). HR fecal metabolomes were enriched in simple sugars, whereas HC contained more nitrogenous compounds. Integrative genome-metabolic modeling of compounds that significantly differentiate asthma-risk groups revealed risk-dependent interactions with community-encoded metabolic potential (CEP), for arabinose and agmatine, whose fecal concentrations are linked with *B. cellulosilyticus* and *H. effluvii* functional traits respectively. These findings suggest that microbial-influenced metabolic differences associate with asthma risk at 6 months, with *B. cellulosilyticus* and *H. effluvii* emerging as candidate bacteria influencing this observed metabolic remodeling.

**Impact statement:** Leveraging a random forest classifier, we identified three bacterial species (*Bacteroides cellulosilyticus, Hungatella effluvii*, and *Enterocloster aldenensis)* as distinguishing features enriched in healthy 6-month old infant microbiomes compared to those at high risk of asthma development (HR). We developed an approach to integrate metabolomics and metagenomic-derived microbiome community encoded potential (CEP) with clinical outcomes to identify fecal metabolites whose concentrations are likely to be influenced by the microbiome. Fecal arabinose concentrations were positively associated with CEP in healthy infants, but not in HR subjects who exhibited elevated concentrations irrespective of CEP. These data implicate microbial activity as a contributor to the concentration of this metabolite in healthy but not HR infants. With a leave-one-out-cross-validation, we identified *B. cellulosilyticus* as a contributor to fecal arabinose concentrations. Our data indicate that microbial functional deficits in HR infants is associated with altered gut metabolic dysfunction during microbiome maturation.

**Data summary:** Durack et. al [1] is the source of the metabolomics data utilized in this study. The authors confirm that all other supporting data, code and protocols have been provided within the article or through supplementary data files.

## 5. Introduction

Atopic diseases, such as allergic asthma, are amongst the most prevalent chronic inflammatory diseases of childhood and are frequently characterized by Th2 inflammation and elevated circulating immunoglobulin E (IgE) concentrations[2]. Although risk alleles associated with atopy and asthma have been identified, the rapid increase in prevalence, particularly in industrialized nations, cannot be explained by host genetics alone[3]. Several studies have linked early life fecal microbiota perturbation and metabolic dysfunction with increased risk of atopy and asthma in childhood[1, 4–6], though little is known of the gut microbiome functional traits that characterize asthma risk, particularly in later infancy.

We previously compared fecal samples collected longitudinally over the first year of life from healthy control (HC) and high-risk for asthma (HR; placebo arm) infants in the Trial of Infant Probiotic Supplementation (TIPS)[1, 7]. HR infants exhibited delayed fecal microbiota diversification and a significantly distinct stool metabolic profile at 6 and 12 months of age[1, 7]. Due to the increased number of significantly differential metabolites at 6 months compared to 12 months[1], the rapid diversification that typically happens at this age when solid food introduction typically occurs[8], and the absence of infant gut metagenomics at 6 months in the current asthma literature, we chose to focus on this critical timepoint. Comparative analysis of asthma-risk groups indicated a significant difference in carbohydrate metabolism characterized by enrichment of simple sugars in the feces of HR compared with HC subjects at 6 months of age. Altered carbohydrate metabolism has also been described in younger (1 month-old) infants at high-risk for atopy, indicating that it represents a consistent feature of those at higher risk of childhood atopy development[5]. However, which microbes contribute to carbohydrate metabolism in the gut microbiome of HR and HC infants is largely unknown. Here we leverage fecal samples collected at 6 months of age from HR and HC infants and apply a multi-omics (genome-resolved metagenomics and metabolomics) approach to characterize the abundance and encoded genomic functions of the infant gut microbiome that differ between HR and HC infants with a specific focus on those fecal microbial species that contribute to altered carbohydrate metabolism.

## 6. Methods

### Study Population

Stool samples for this study were obtained from the Trial of Infant Probiotic Supplementation (TIPS – ClincialTrials.gov ID: NCT00113659) clinical trial and Development of Infant Microbial Evolution (DIMES) birth cohort. Recruitment for TIPS was carried out in the San Francisco Bay Area and consisted of infants at high-risk of childhood asthma development based on an asthma diagnosis of one or both parents.[7] Infants were randomized in this double blind placebo controlled trial of daily oral *Lactobacillus rhamnosus* GG supplementation, and stool samples were collected at birth, 1, 3, 6, 9, and 12-months of age. Stool samples collected at 6 months of age from placebo treated infants in the TIPS trial served as our high-risk (HR) population for this study. DIMES also recruited healthy infants (e.g., no history of asthma, eczema or rhinitis) from the Bay Area. DIMES stool samples collected at 6 months of age served as our healthy control (HC) population for this study. This pilot study focused on fecal samples collected at 6-months of age because of paired extant untargeted metabolomics data available for many of these samples[1], the increased number of significantly differential metabolites at this timepoint compared to 12 months[1], and the rapid diversification that typically happens at this critical age when solid food introduction typically occurs[8]. A total of 15 HC and 11 HR samples had available metabolomics for analysis, and as some samples had been fully consumed in earlier experiments, genome resolved metagenomic analysis could only be conducted on 12 HC and 7 HR samples. Across all samples evaluated in this work, 10 HC and 7 HR possessed paired metagenomic and metabolomic datasets (61% of samples).

### DNA Extraction

DNA was extracted from 6-month old infant stool samples. Frozen samples were maintained on dry ice while a 4 mm punch biopsy with plunger (VWR International) was used to aliquot 0.3 grams of sample into a Lysing Matrix E tube (LME; MP Biomedicals) with 500 µL hexadecyltrimethylammonium bromide (CTAB, Sigma-Aldrich) and 500 µl phenol:chloroform:isoamyl-alcohol (25:24:1, Sigma-Aldrich). Samples were incubated at 65 ºC for 15 minutes. Following incubation, samples were homogenized in a Prep-24 homogenizer (MP Biomedicals) at 5.5 m/s for 30 seconds and centrifuged at 16,000 x g for 5 minutes at 4 ºC. The aqueous phase was transferred to sterile 1.5 mL tubes. An additional 500 µL of CTAB buffer was added to the LME tubes and incubation, homogenization, and centrifugation steps were repeated. The aqueous phases from both extractions were combined in a sterile 1.5 mL tube. An equal volume of chloroform was mixed with each sample, followed by centrifugation at 16,000 x g for 10 minutes at 4 ºC. The aqueous phase (600 µL) was transferred to a clean tube, combined with 2 volumes (1,200 µL) of polyethylene glycol (PEG, Sigma-Aldrich) and stored overnight at 4 ºC to precipitate DNA. Microcentrifuge tubes were centrifuged for 10 min at 16,000 x g, prior to washing the resulting DNA pellets with 300 µL of ice-cold 70% ethanol. Pellets were air-dried for 10 minutes and re-suspended in 100 µL of sterile water. DNA from stool samples was quantified using the Qubit dsDNA Broad Range Assay Kit, diluted to 10 ng µL^-1^

### Metagenomic Sequencing

Purified total DNA from the extracted 12 HC and 7 HR infant samples was sent to the Vincent J. Coates Genomic Sequencing Laboratory at the California institute for Quantitative Biosciences. DNA was fragmented and 150 bp x 2 (paired-end) libraries were constructed and sequenced on an Illumina HiSeq 4000 instrument to an average depth of 10M total reads per sample. (www.qb3.berkeley.edu/gsl).

### Metabolomics

Extant untargeted metabolomics data previously generated in Durack *et al*[1] was used for this study. Briefly, 200 mg of stool samples from patients collected at 6 months (n=15 HC, n=11 HR) underwent ultrahigh performance liquid chromatography/tandem mass spectrometry (UPLC–MS/MS) and gas chromatography–mass spectrometry (GC–MS) by Metabolon (Durham, NC), using their standard protocol (http://www.metabolon.com/). Compounds were compared to Metabolon’s in-house library of purified standards, which includes more than 3,300 commercially available compounds.

### Metagenomics Analysis

#### Read Processing, Assembly and Binning

Raw metagenomic reads were processed using a Snakemake-based workflow. In brief, reads were first assessed using FastQC (v0.11.09)[9], followed by adapter removal and quality trimming to a mean of Q20 using BBDuk (v39.10)[10] and Sickle (v1.33)[11]. Assemblies for individual samples were generated with IDBA-UD (v1.1.3, r252)[12] using a k-mer range of 20-150 nucleotides. To improve contig recovery, three merged read sets constructed by concatenating all reads from (i) HC samples, (ii) HR samples, and (iii) all samples combined. Co-assemblies of these merged read sets were generated with IDBA-UD (v1.1.3, r252)[12] using a k-mer range of 30-150 nucleotides. A larger minimum k-mer was set for concatenated assemblies due to memory requirements. Following assembly, contigs < 1 kb were discarded. COBRA (v1.2.3)[13] was run on assembled contigs ≥ 5 kb to detect circular elements and extend contigs. Coverage of assembled contigs was evaluated by mapping sample reads back to contigs using Bowtie2 (v2.5.2) [14].

Assemblies were independently binned using an automated pipeline integrating MetaBAT2 (v2.12.1)[15], MaxBin2 (v2.2.7) [15, 16], CONCOCT (v1.1.0)[17], and VAMB (v3.0.9)[18]. Contigs ≥ 2.5 kb were retained for binning; coverage of contigs in each sample vs all other samples was calculated by cross-mapping reads using SNAP (v2.0.1)[19] and a filtered coverage table was produced using jgi_summarize_bam_contig_depths from the MetaBAT2 package. The best bins (MAGs) across all 4 binning methods in each sample were selected using DAS Tool[20] (v1.1.1; score ≥0.3).

#### Genome Aggregation, Coverage Profiling and Functional Annotation

MAGs from all samples were aggregated, quality-checked (CheckM v1.2.1; ≥60% completeness, ≤10% contamination)[21], and dereplicated with dRep (v3.4; ≥95% ANI, ≥10% overlap)[22]. Taxonomy was assigned with GTDB-Tk (v2.1.1, r207)[23]. Genetic codes were inferred with Codetta (v2.0)[24], ORFs predicted with Prodigal (v2.6.3)[25], and tRNAs with tRNAscan-SE (v2.0.12)[26]. The final genome set consisted of 82 species-representative MAGs with no MAG sharing more than 95% average nucleotide identity (ANI) to any other representative MAG. This set of MAGs was then used as a reference database for microbial abundance estimation. Predicted proteins were annotated using KofamScan (v1.3.0)[27], and only hits passing the adaptive thresholds were retained to build KO-based functional profiles.

To assess microbial abundance and composition in each sample, reads from all samples were mapped to the species-representative MAGs database using Bowtie2 (v2.5.2)[14] to generate per-sample alignment files. Coverage statistics were then computed using CoverM (v0.7)[28], which summarized genome-level coverage and read counts within each sample.

All statistical analysis were performed in R (v4.4.2) and visualizations generated using ggplot2 (v3.5.1)[29]. We initially removed MAGs from our analysis that were present in ≤ 2 samples (n=20 MAGs) resulting in 62 MAGs being retained for analysis. Read counts were zero imputed with zero multiplicative replacement and normalized using center log-ratio transformation. Beta diversity was assessed using Aitchison distances calculated from CLR-transformed data. To test for differences in overall community composition between asthma risk groups, we performed a PERMANOVA (permutational multivariate analysis of variance) using the adonis2 function from the vegan R package (v2.7.1)[30], with 999 permutations.

#### Feature Selection, Co-occurrence, and Functional Enrichment

To identify microbes associated with asthma risk group classification, we applied the Boruta algorithm, a wrapper feature selection method built around Random Forests, using the Boruta R package (v8.0.0)[31]. Feature importance scores were computed by comparing the relevance of each real feature to that of permuted shadow features across multiple randomized forests. To ensure stability of feature selection, Boruta was run across a grid of random seeds (n = 3: 11, 145, 224) and different numbers of decision trees (ntree = 1000 - 5000, in steps of 500), with a maximum of 200 iterations per run (maxRuns = 200). For each run, importance scores were extracted for all features and normalized as z-scores within each iteration. These z-scores were then aggregated across all seeds and tree counts into a unified dataset for visualization. Genomes confirmed to distinguish asthma risk group underwent a Wilcoxon rank-sum test, and p-values were adjusted for multiple testing using the Benjamini-Hochberg method. To account for class imbalance, Boruta was repeated using class-weighted Random Forests, with weights inversely proportional to class frequency implemented via the “class.weights” parameter.

Microbial co-occurrence analysis was performed using the cooccur R package (v1.3)[32]. A binary presence/absence matrix was generated by thresholding CLR-transformed genome abundances (presence defined as >0). Co-occurrence was assessed using a probabilistic null model (type=“spp_site”, thresh=TRUE, spp_names=TRUE), with species pairs having an expected co-occurrence <1 excluded from analysis. Of 1,891 possible species pairs, 700 (37.0%) were filtered, and 1,191 were tested for non-random associations. Pairs were classified as positive, negative, or random based on the statistical significance of their observed vs. expected co-occurrence (p < 0.05).

Differential enrichment of KEGG Orthologs (KOs) between *Bacteroides cellulosilyticus* and the other 61 genomes evaluated was assessed using Fisher’s Exact Test. KO count tables were filtered to exclude KOs observed in fewer than 10 genomes. *Bacteroides* KO counts were summed separately from background genomes, and 2×2 contingency tables were constructed for each KO. Fisher’s Exact Test was applied to each table to test for significant differences in KO presence between groups, and p-values were adjusted for multiple testing using the Benjamini-Hochberg method.

### Metabolomics Analysis

Of the 692 compounds detected in the untargeted metabolomics, 571 were present in at least half of the samples of each group (HC and HR) and retained for analysis. Metabolite intensities were normalized using probabilistic quotient normalization to correct for sample dilution, followed by log_2_ transformation. Missing values were imputed within phenotype groups using the quantile regression imputation of left-censored data (QRILC) algorithm under a left-censoring assumption. To control for batch effects while preserving biological variation associated with asthma risk, ComBat from the sva package (v3.54.0)[33] was applied using run day as the batch variable and the group (HC or HR) as a covariate in the design matrix.

Principal component analysis (PCA) was performed on the log_2_-transformed, batch-corrected data to visualize overall variance and group separation. The contribution of asthma-risk status to global metabolic variation was evaluated using PERMANOVA (adonis2, 9,999 permutations) on Euclidean distances derived from scaled metabolite intensities. Euclidean distances were chosen because scaling produces continuous, non-compositional features, and keeps the analysis similar to Aitchison distance metrics used in the metagenomic analysis.

Orthogonal Partial Least Squares Discriminant Analysis (OPLS-DA) was performed using the ropls R package (v2.38.0)[34] to identify metabolites that discriminate between HR and HC infant groups. Model performance was evaluated with 5-fold cross-validation and 9,999 label permutations (permI = 9999). The top 100 metabolites ranked by variable importance in projection (VIP) scores from the OPLS-DA model were tested for differential abundance between HR and HC groups using limma (v3.62.2)[35]. Limma fits metabolite-wise linear models and employs empirical Bayes moderation to stabilize variance estimates across features. Resulting p-values were adjusted for multiple testing using the Benjamini–Hochberg false discovery rate (FDR).

### Metagenomic-Metabolomic Integrative Community Encoded Metabolic Potential (CEP)

Of the top 100 metabolites ranked by VIP, 60 had an assigned KEGG compound identifier. Among these, 39 were significantly different between HC and HR infants as tested via Limma described above. 33 of these 39 significant compounds were linked to KOs represented in the metagenomic annotations. Of these 33 compounds, 26 had KEGG IDs which were present within the genomes analyzed in this study. For these 26 compounds, a genome-by-compound (GxC) weight matrix was generated by counting, per genome, the number of KO annotations linked to each compound. A genome-by-sample (GxS) matrix was generated by converting the CLR-normalized genome abundance table to sample-wise proportions by exponentiation and then rescaling each sample so that all genomes within that sample sum to 1. CEP was computed as the matrix product of GxS and GxC to cancel out genomes producing a samples-by-compound (SxC) matrix of community-encoded metabolic potential. The z-score of CEP was calculated across samples for standardization then fit to a robust rank-based regression (rfit) of measured metabolite abundance on CEP with asthma risk-group as an interaction below:

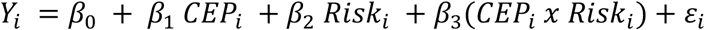

*Y*_*i*_ is the measured abundance for compound i. Risk is a binary indicator for the infant group (0= healthy, 1=high-risk). Each coefficient corresponds to a specific biological interpretation: *β*_0_ is the intercept (expected metabolite abundance for healthy infants when CEP = 0), *β*_1_ captures the CEP-metabolite association in healthy infants, *β*_2_ captures baseline differences between risk groups, and *β*_3_ tests whether the CEP-metabolite relationship differs by asthma-risk group. *ε*_*i*_ is the error term. For each compound fit to the model above, the interaction effect (*β*_3_) was FDR-adjusted using Benjamini-Hochberg. Given our small sample set, p_int_ CEPxRisk values <0.1 prior to FDR correction were investigated. To estimate the predictive influence of the individual genomes on the compound model, a leave-one-out-cross-validation (LOOCV) of the robust model using the full CEP predictor versus a CEP predictor missing the genome of interest was calculated and defined as Δr^2^_cv_. Genome and compound-level summaries were computed from the positive Δr^2^_cv_ values.

## 7. Results

### *Bacteroides cellulosilyticus, Hungatella effluvii*, and *Enterocloster aldenensis* distinguish asthma risk groups

Principal coordinates analysis (PCoA) based on an Aitchison distance analysis of species-level genome counts did not identify significant microbiome species differences between the two risk groups (**Figure 1A**). However, using a permutative random forest algorithm for feature selection (boruta), several genomes significantly distinguished asthma risk groups (**Figure 1B**). Comparative analysis of the relative abundance of the genomes which distinguished the asthma group greater than the median of the shadow max identified three that exhibited a significant difference (p < 0.01) between risk groups; *Bacteroides cellulosilyticus, Hungatella effluvii*, and *Enterocloster aldenensis*. A class-weighted secondary analysis was highly concordant with 99.7% of the random forest decisions agreeing with the unweighted analysis indicating these results are robust to class imbalance (**Figure 1C**). All three genomes are significantly more abundant in HC infant microbiomes (FDR p=0.012; **Figure 1D**). While no significant co-occurrence was detected between *B. cellulosilyticus* and *H. effluvii* or *E. aldenensis* across samples (**Figure 1E**), a positive association between *H. effluvii* and *E. aldenensis* was observed. This suggests that *B. cellulosilyticus* differentiates asthma risk groups independently of other species, whereas *H. effluvii* and *E. aldenensis* form a co-occurring pair with a common association pattern.

**Fig. 1.**
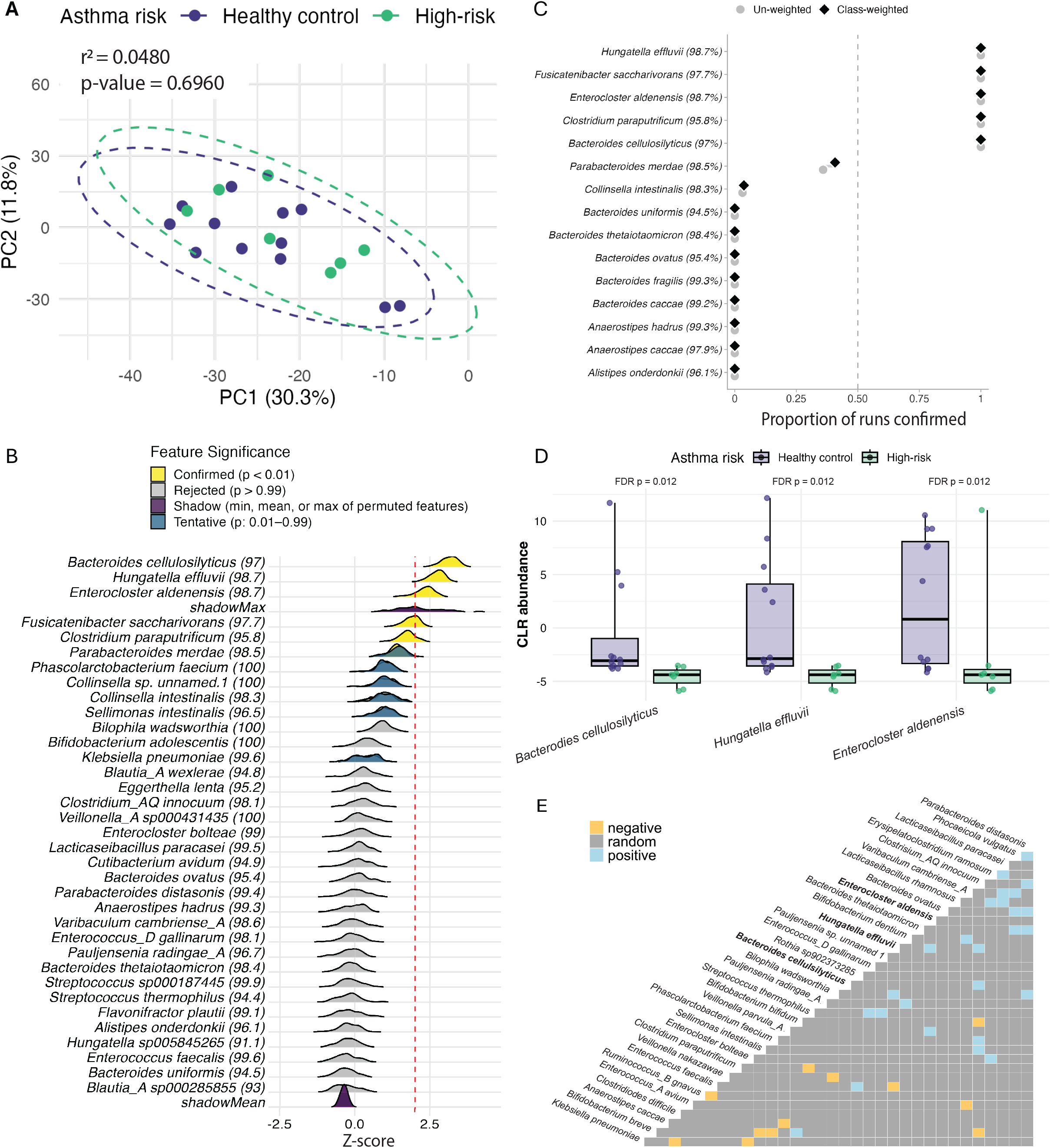
*Bacteroides cellulosilyticus, Hungatella effluvii* and *Enterocloster Aldenensis* are present at higher abundances in 6 month-old infants at low-risk for asthma. A) PCoA of Aitchison distances of CLR-transformed species-level genome abundances. B) Z-scores of mean feature importance to asthma risk status across multiple Boruta runs with varying random seeds and numbers of trees (1,000–5,000) for each microbial species. Features are ranked by median importance and colored by their final Boruta decision. C) Proportion of runs confirming the genome as a distinguishing feature between the un-weighted and weighted Boruta analysis. D) CLR abundance of *Bacteroides cellulosilyticus, Hungatella effluvii*, and *Enterocloster aldenensis* by asthma risk status. E) Pairwise co-occurrence matrix of genomes. Colors indicate positive, negative, and non-significant associations.

### Fecal metabolomes are distinct in HR and HC infants

Unlike the species-resolved metagenomic data, fecal metabolomic profiles were significantly different in HR and HC infants (PERMANOVA; r^2^ =0.068, p=0.015), suggesting broad metabolic differences across these groups (**Figure 2A**). OPLS-DA identified the top 100 metabolites by variable importance projection (VIP) for subsequent statistical testing (**Figure 2B**). Sixty metabolites were found to be differentially abundant between HR and HC samples (FDR < 0.05; |log_2_FC| > 1) falling into the following classes: lipids (20), nitrogenous compounds (12), carbohydrates (11), xenobiotics (12), cofactors and vitamins (3), and peptides (2). All 11 differentially abundant carbohydrates were significantly enriched in HR infant samples and 9 of 12 nitrogenous compounds were significantly enriched in the HC infant samples (**Figure 2C**).

**Fig. 2.**
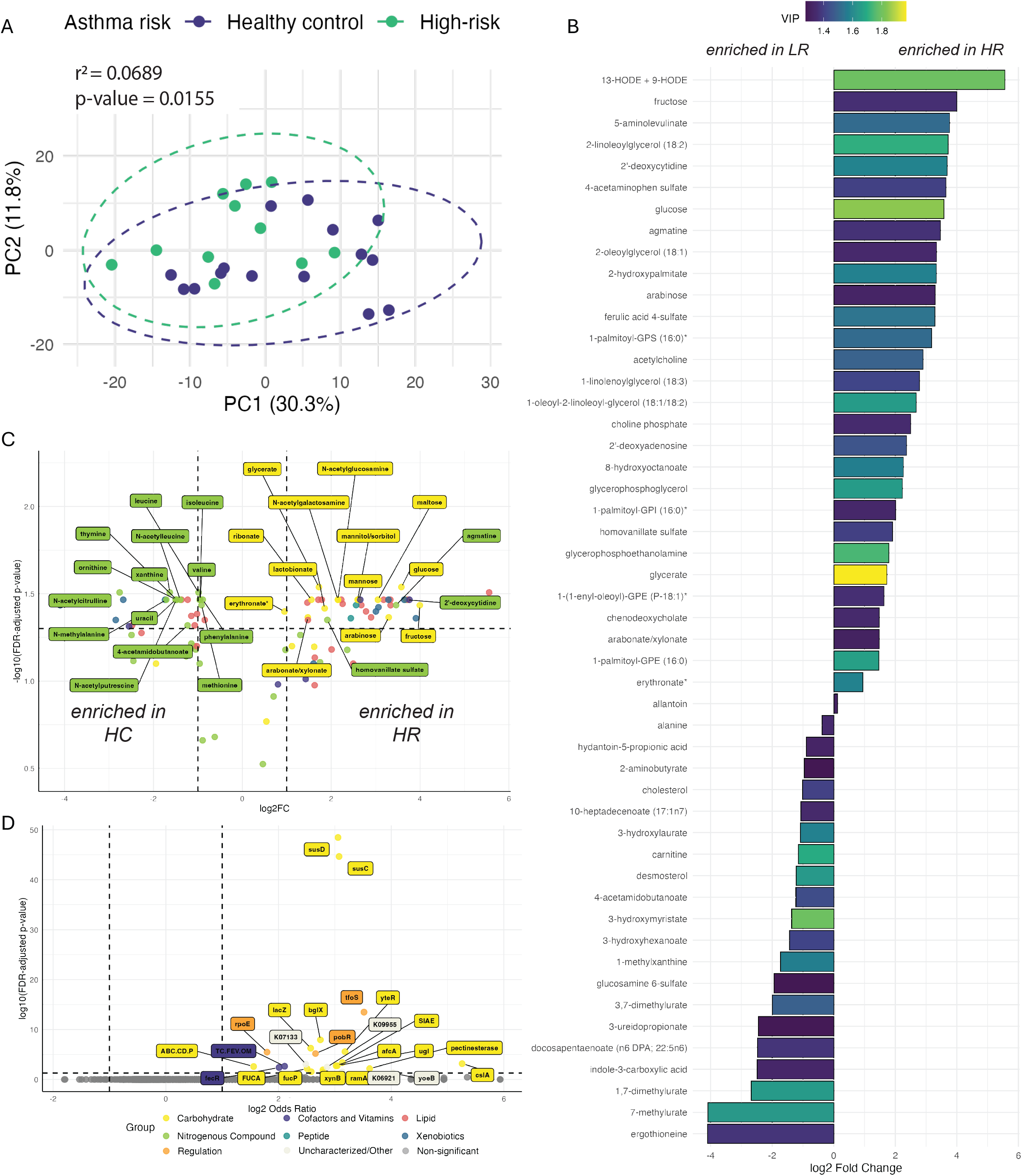
Infants at high-risk for asthma have a stool metabolome characterized by increased simple carbohydrates, and the *Bacteroides cellulosilyticus* genome is enriched for carbohydrate-related genes. A) PCA of auto-scaled log-normalized stool metabolomics data. B) Top 50 metabolites predictive of asthma risk status colored by Variable Importance in Projection (VIP). C) Statistical significance and log-fold change of the top 100 compounds identified by OPLS-DA. Carbohydrates and nitrogenous compounds with significantly differential abundance (ANOVA FDR-adjusted P-value < 0.05) are labeled. D) Genes significantly enriched in the *Bacteroides cellulosilyticus* genome compared to the 61 other species-level genomes evaluated in this study.

### *Bacteroides cellulosilyticus* and *Hungatella effluvii* influence distinct metabolite-CEP interactions based on asthma-risk

We next focused our analysis on the functional potential within the fecal microbiome which, we hypothesized, influences the infant gut metabolome. Glycan degradation genes (*FUCA, xynB*, and *bglX)* and glycan import genes (*SusC and SusD*) were significantly more abundant (FDR < 0.05) in *B. cellulosilyticus* genomes compared to all other MAGs in this analysis (**Figure 2D**). To better understand which fecal metabolites may be influenced by these microbes we developed and deployed an integrative metagenomic-metabolomic analytical approach. Community encoded metabolic potential (CEP) for a particular metabolite was calculated from KO function abundance data (stratified by genome) and correlated with the observed metabolite abundance. We performed this analysis on the subset of data with paired metagenomic and metabolomic samples (HR=7, HC=10). Of the 26 metabolites assessed, interactions between agmatine (p_int_ = 0.0529) and arabinose (p_int_ = 0.0598) with asthma-risk passed the p_int_ significance threshold, however none of these findings remained significant following multiple testing correction (p_int.adj_ = 0.6539 for all; **Figure 3A-D**).

**Fig. 3.**
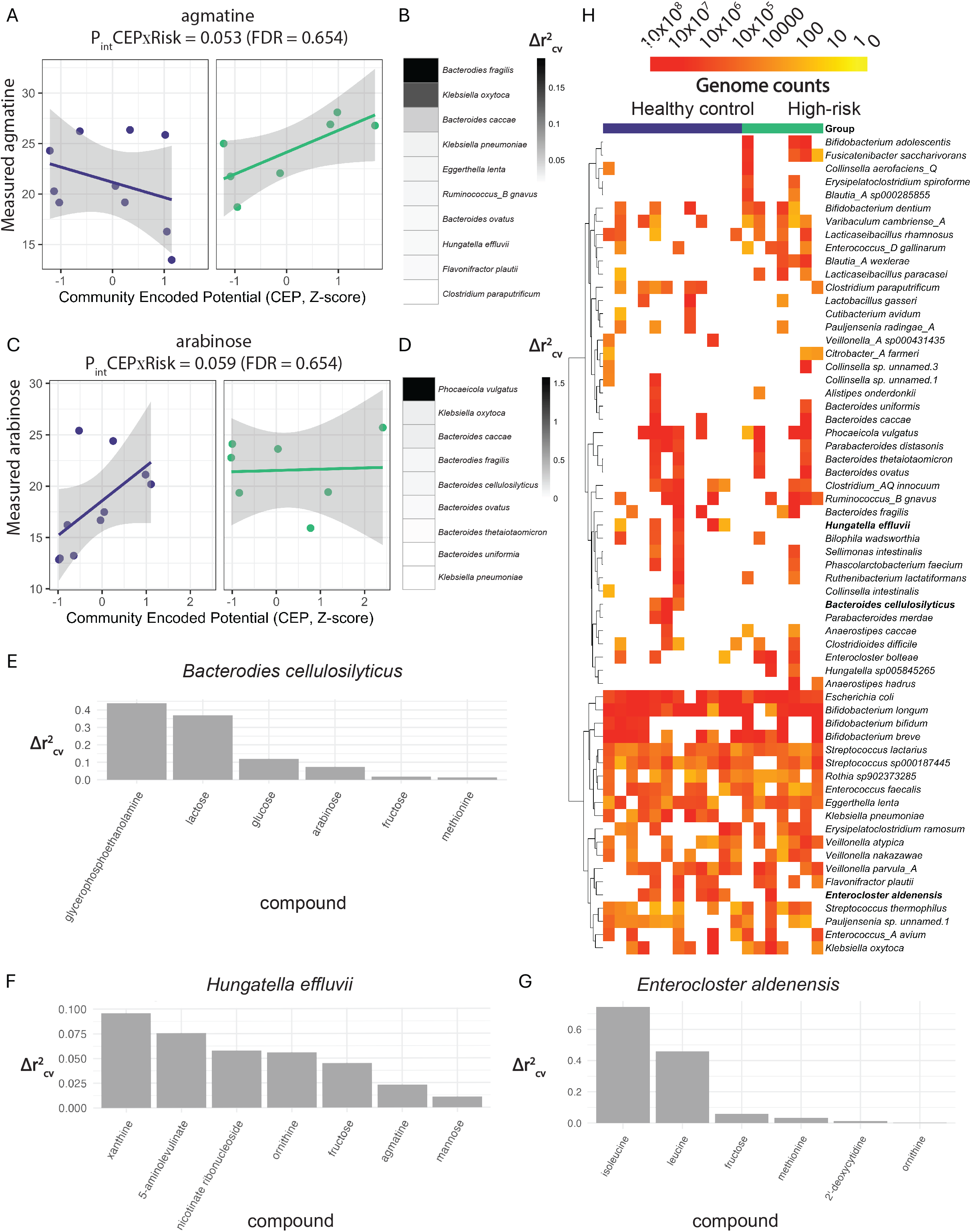
Risk-dependent metabolite–community encoded metabolic potential interactions reveal *B. cellulosilyticus* and *H. effluvii* as contributors to arabinose and agmatine abundance. A,C) Robust regression models evaluating the interaction between community-encoded metabolic potential (CEP) and asthma-risk group for agmatine (A) and arabinose (C). B,D) Shaded regions denote standard error of the regression. Leave-one-genome-out Δr^2^_cv_ analyses identifying genomes which influence the CEP–metabolite models for agmatine (B) and arabinose (D). E-G) Compound-specific Δr^2^_cv)_ profiles for *B. cellulosilyticus* (F), *H. effluvii* (G), and *E. aldenensis* (H). H) Log-transformed genome counts for all 62 species-level MAGs across healthy control (HC) and high-risk (HR) infant stool samples (n = 19).

Further analysis of these relationships indicated that agmatine is negatively associated with CEP in HC samples but positively associated with CEP in HR samples (**Figure 3A)**. Fecal arabinose concentrations were found to be positively associated with CEP in HC samples implicating microbial activity as a key contributor to the concentration of this metabolite in these infants. Notably, we found that arabinose is maintained at a relatively higher concentration in HR infants regardless of CEP, implicating dietary sources of this sugar in high-risk infants (**Figure 3C)**.

The primary contributors to the CEP x Compound models for both agmatine and arabinose are organisms that are present in both asthma risk groups (*B. fragilis* and *P. vulgatis*; **Figure 3B,D,H**). However, both *H. effluvii and B. cellulosilyticus*, rank highly amongst the genomes that contribute to the variance in fecal agmatine and arabinose concentrations. *H. effluvii* is the 8th most important contributor to agmatine x CEP model variance and *B. cellulosilyticus* is the 5th most important contributor to arabinose x CEP model variance (**Figure 3B, D**). This implies that despite a small sample size and uneven distribution of these organisms, they are important contributors to the abundance of these fecal metabolites, particularly in healthy infants (**Figure 3A-D;H)**. Furthermore, we specifically evaluated which compound’s abundance and CEP variance was most explained by the genomic capacity of *B. cellulosilyticus, H. effluvii*, and *E. aldenensis. B. cellulosilyticus* primarily explained carbohydrate variance (glycerolphosphoenthanolamine, lactose, glucose, arabinose, fructose; **Figure 3E**), while *H. effluvii* and *E. aldenensis* associated with nitrogenous compounds (**Figure 3F -** *H. effluvii*: xanthine, 5-aminovulinate, nicotinate ribonucleoside, ornithine, and agmatine; **Figure 3G -** *E. aldenensis*: isoleucine and leucine).

## 8. Figures and tables

## 9. Discussion

The principal factors influencing infant gut microbial community structure at 6 months of age are dietary exposures, including breastfeeding, formula feeding, and the introduction of solid foods[8]. These dominant influences may mask more subtle ecological or functional differences associated with early-life risk factors for childhood asthma development. Consequently, large-scale shifts in overall microbiome composition are not expected to distinguish asthma risk groups, a pattern observed both in our study and in prior investigations[1, 36]. Recognizing this, our study aimed to detect more nuanced early-life microbial determinants or correlates of asthma risk. This approach acknowledges that early-life microbial risk factors may operate through discrete microbes, metabolic outputs, or host-microbe interactions, rather than broad community-level restructuring.

Using a feature-selection approach to identify the organisms that most strongly differentiated HR and HC samples, we identified three microbes (*Bacteroides cellulosilyticus, Hungatella effluvii, and Enterocloster aldenensis)* which were associated with asthma risk and enriched in those infants at lower-risk suggesting they may play a role in asthma protection. *B. cellulosilyticus* has been reported to have anti-inflammatory effects in a gnotobiotic colitis mouse model reducing IL-6, TNF-α, and PI3K/Akt phosphorylation[37]. Furthermore, *B. cellulosilyticus* is capable of utilizing a wide array of carbohydrates including glucose, arabinose, N-acetylgalactosamine, N-acetylglucosamine, mannose, and fructose[38], all of which were depleted in the stool metabolome of HC compared with HR infants in our study. Our integrative analysis of metabolomic data and CEP also points to *B. cellulosilyticus* as an important contributor to fecal carbohydrate relative abundances, as its largest related influences are on carbohydrate metabolites. More broadly, we observed the CEP had a positive association with arabinose abundance in low-risk infants and minimal association with the relative abundance of this sugar in those at high-risk. This suggests that the gut microbiome may differentially processes substrates in a health status-specific context, and that factors not captured in this study such as dietary differences may play a significant role in dictating these microbial interactions. *B. cellulosilyticus, H. effluvii*, and *E. aldenensis* were depleted in HR infant fecal microbiomes. *B. cellulosilyticus* is enriched for genes encoding enzymes that utilize the specific sugars found in higher concentrations in HR infants, indicating that the loss of these species and their encoded functional traits may contribute to the excess sugars observed in the feces of these infants.

*H. effluvii* and *E. aldenensis* are less well-studied organisms, and the associations between these species and nitrogenous compounds suggests may be linked to the variation in these compounds, several of which have known roles in inflammation[39]. Agmatine, which was relatively enriched in the HR infant stool metabolome is known to be produced by commensal organisms and has been implicated in inflammatory processes[39]. While the precise impact of agmatine on early-life immune development remains unclear, our findings pinpoint *H. effluvii* and *E. aldenensis* as contributors to nitrogenous compound metabolism in the gut microbiome of HC infants. Our data suggests that loss of these species is associated with increased concentrations of agmatine in the gut of infants at higher risk of childhood asthma development.

While this study moves beyond exclusively evaluating taxonomic or metabolite-level associations with asthma risk, and points to *B. cellulosilyticus, H. effluvii*, and *E. aldenensis* as organisms associated with lower asthma risk, several limitations should be noted. Due to utilizing previously processed samples, the sample size and sample overlap between metagenomic and metabolomic datasets was limited. Sample size limitations become apparent in CEP modeling where a large number of features and co-variates are evaluated and findings fail to pass our FDR threshold. Furthermore, the current annotations available in the KEGG database, which was used for both gene enrichment analysis and CEP modeling, poorly cover pathways such as microbial lipid metabolism that are known to be important in mediating inflammation and asthma risk[40, 41]. Future work should include a significantly larger sample size of paired metagenomic and metabolomic datasets. Also, evaluating clinical incidence of disease rather than disease risk based on parental presence of asthma would be preferable. Nonetheless, this proof-of-concept study identifies specific bacteria that both have the encoded potential to interact with the concentrations of several carbohydrate and nitrogenous compounds that differentiate HR and HC infants. Finally, our results from CEP modeling indicate that more complex interactions between encoded microbial metabolic potential and metabolite abundances exist, suggesting that additional factors such as diet and host metabolic capacity may influence metabolite concentrations, particularly in those at higher disease risk.

## 10. Author statements

### 10.1 Author contributions

HMS: Formal analysis; Visualization; Software; Writing – original draft; Writing – review & editing.

CEIA: Formal analysis; Validation; Visualization; Writing – review & editing.

ARP: Conceptualization; Investigation; Writing – original draft.

JD: Investigation.

MM: Project administration.

MDC: Data curation; Project administration; Resources.

SD: Funding acquisition; Software; Supervision; Writing – review & editing.

SVL: Conceptualization; Funding acquisition; Supervision; Writing – review & editing.

### 10.4 Ethical approval

The Trial of Infant Probiotic Supplementation (TIPS – ClincialTrials.gov ID: NCT00113659) clinical trial and Development of Infant Microbial Evolution (DIMES) birth cohort were both approved by the University of San Francisco, California’s IRB #2019-10582.

